## Supplementary material for "Conservation of the *Toxoplasma* conoid complex proteome reveals a cryptic conoid in *Plasmodium* that differentiates between blood- and vector-stage zoites": Gel and blot raw images

S1_Raw_Images supporting gels and blots

Western blot images used for Fig 2: A) chemiluminescent signal for streptavidin-detected biotinylated proteins; B) gel image showing prestained molecular weight markers (M); C) merge of inverted image A on B. Lane numbers; 1, paraental; 2, SAS6L-BirA*; 3, RNG2-BirA*; 4, MORN3-BirA*.

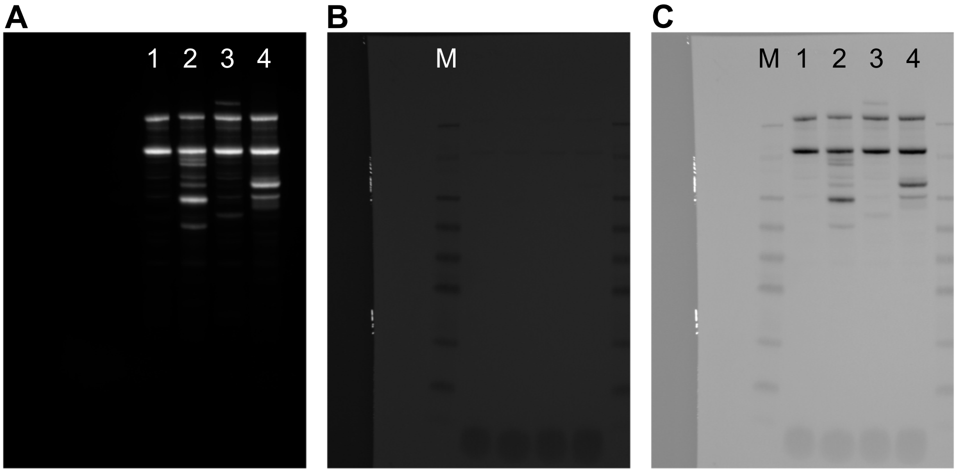


DNA gels used in Fig S9. Gene number is indicated with four corresponding lanes below: Lanes 1 (wildtype control) and 2 (tagged cell line) show PCR results for gene-specific GFP integration, lanes 3 (wildtype control) and 4 (tagged cell line) show positive control PCR. X, indicates lanes not included in Fig S9, and all unboxed lanes are unrelated to this report.

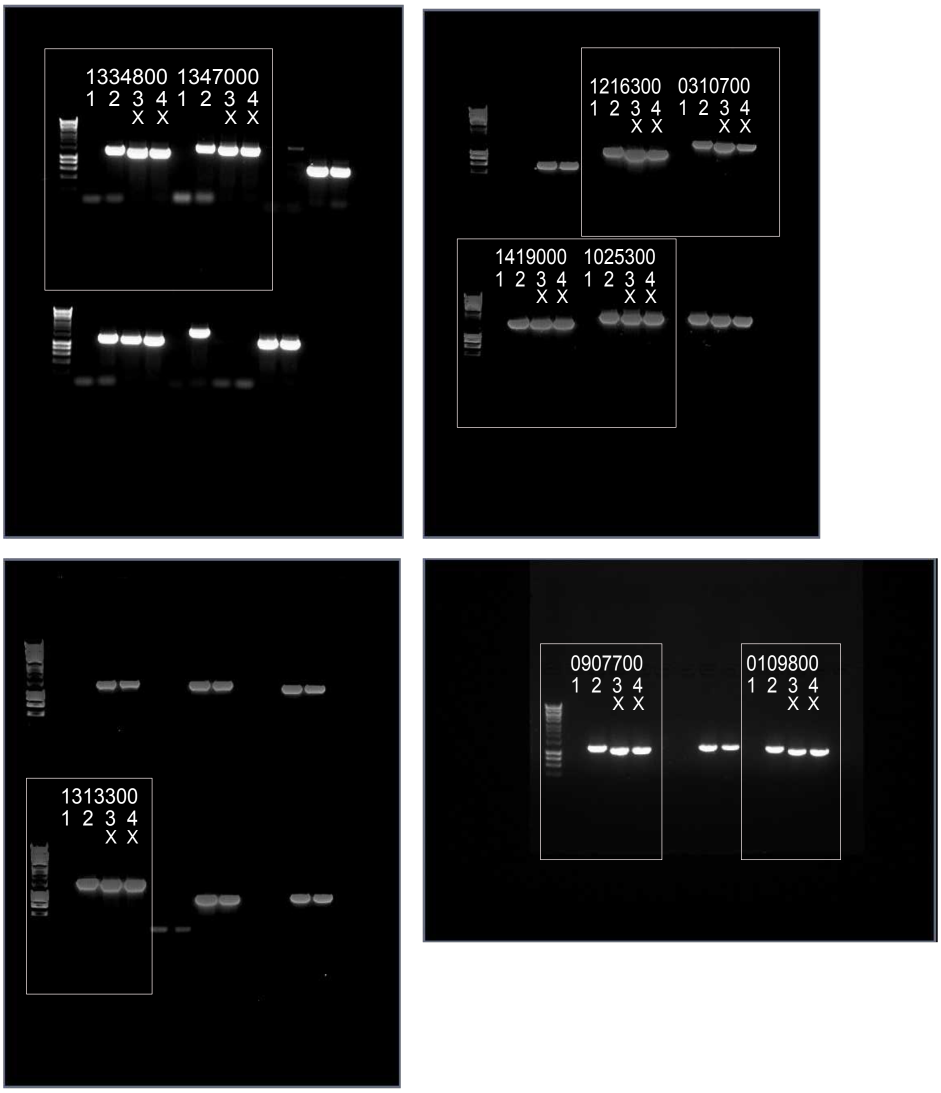
