## Supplementary figures and images for "Conservation of the *Toxoplasma* conoid complex proteome reveals a cryptic conoid in *Plasmodium* that differentiates between blood- and vector-stage zoites"

### Figure S5

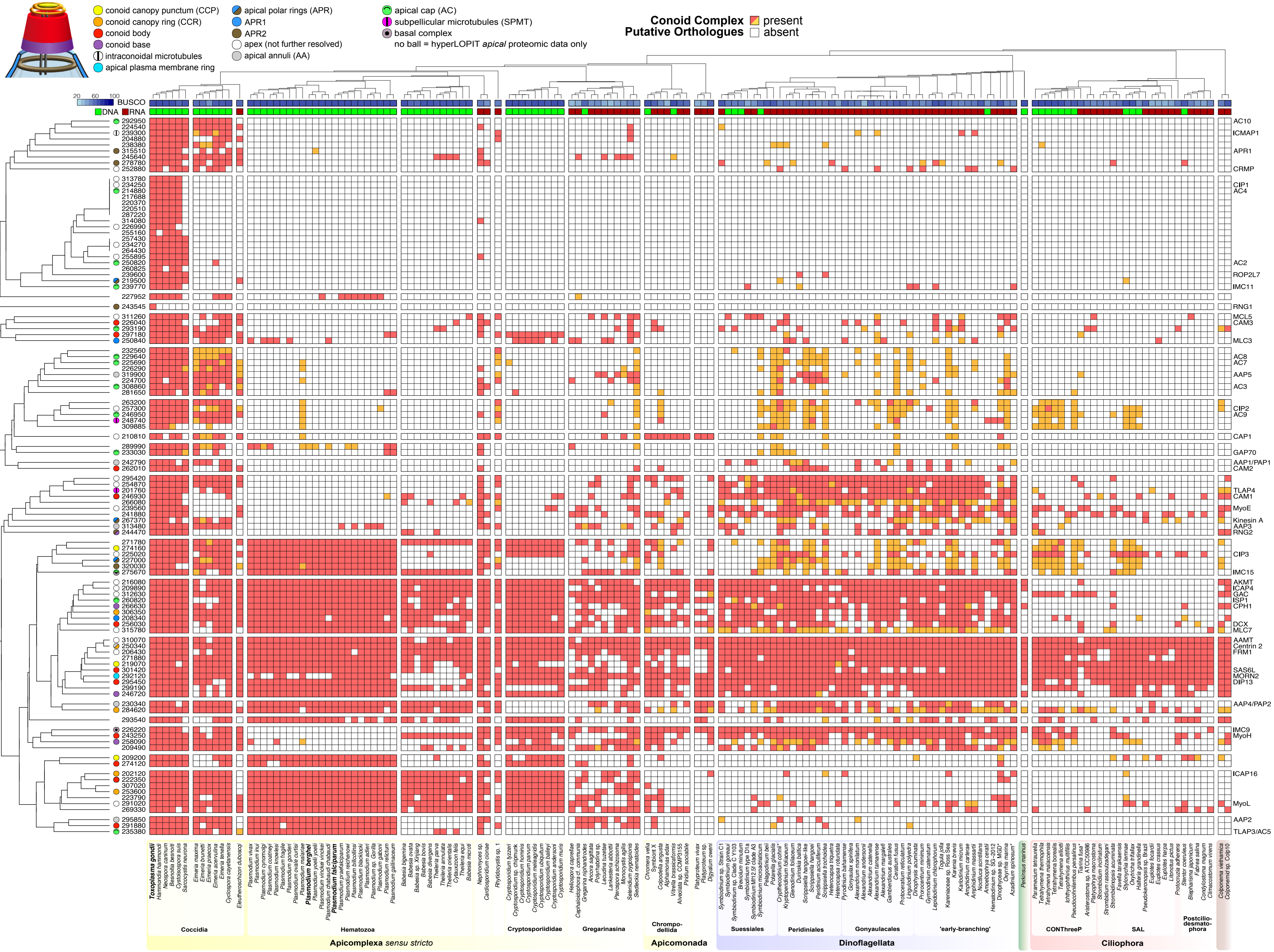
